# Knowledge and self-care practice of leprosy patients at ALERT Hospital, Ethiopia

**DOI:** 10.1101/378067

**Authors:** Lisanawork Girma, Kidist Bobosha, Tsegaye Hailu, Edessa Negera

## Abstract

**Background:** Leprosy is a chronic infectious disease of public health importance and one of the leading causes of permanent physical disability. The integration of leprosy service to the general health system in Ethiopia made leprosy patients to be seen by non-leprosy specialists which could lead to misdiagnosis and delay in treatment. In addition to the multi-drug treatment, patient self - care practice is crucial for the successful treatment of the disease. This study was aimed to assess the knowledge and self-care practices of leprosy patients and associated factors at ALERT leprosy referral Hospital in Ethiopia.

**Method:** A cross-sectional study was conducted ALERT leprosy referral Hospital, Addis Ababa, Ethiopia. A total of 424 leprosy patients were interviewed using pre-tested structured questionnaires. The questionnaires included core points such as socio-demographic characteristics, knowledge of leprosy and self-care practices. Bloom’s cut off point was used to describe the knowledge and self-care practice of the respondents and statistical significance was assessed at 95% confidence interval with 5% of level of significance.

**Result:** The knowledge score of the respondents was poor for 276 (65.1%) and good for 148 (34.9%). The level of knowledge was significantly varied within age groups (P= 000), sex (P=000), marital status (p=0.003), educational status (p=000) and income (p-000). More than three-fourth (77.4%) of interviewed patients had poor self-care practice and only less than a quarter (22.6%) of patients had good self-care practice score (P=000). Age (p=0.002), Previous disability due to leprosy (P=000), Knowledge of leprosy (p=0.038) and income (P=0.028) significantly associated with poor self-care practice. On the other hand, educational status, sex, marital status and health education did not associated with leprosy self-care practice.

**Conclusion:** Although, leprosy treatment, disability prevention and rehabilitation program run in the country under general public health care service for decades, poor leprosy self-care practice and poor leprosy knowledge had been confirmed in this study. Therefore, the leprosy service program should re-visit its strategy and mode of delivery to improve the leprosy knowledge and leprosy self-care practices of patients.

**Author summary:** Leprosy is an infectious bacterial disease. It is a curable disease if treated early before disability occurs with the correct regimen. However, treatment cannot reverse disability once occurred. in addition to chemotherapeutic treatment, lack of social and psychological treatment may expose patients to disability as they may not adhere to treatment. To prevent disability it is vital to empower leprosy patients through promoting self-confidence, providing knowledge and self-care skills. In the present study, we investigated the knowledge and self-care practice of 424 leprosy patients at ALERT hospital through face-to face guided interview. It was found that majority of patients had poor knowledge about leprosy. Some factors such as sex, income status, age and educational status of the patient significantly affected the level of leprosy knowledge. Similarly, it was found that 77% of patients had poor self-care practice which significantly varied with income status, knowledge of leprosy, age and having previous disability. Therefore, it is very important to improve the leprosy knowledge and self-care skills of patients. This can be achieved through dealing with issues such as the physical, emotional, intellectual and social aspects of the patients in addition to the chemotherapeutic treatment.

## Introduction

According to World Health Organization (WHO), the prevalence of leprosy has reduced by 90% from 21.1 per 10,000 populations in 1985 to less than 1 per 10,000 populations by 2000 [1]. The prevalence of leprosy has decreased by 97 % in the past 3 decades [2]. It has dropped from 5.2 million in 1985 to 171, 948 in 2017 [2]. On the other hand, new case detection has only dropped by less than 10% in a decade. In 2007, a total 258,133 new cases were reported to WHO and 214,783 new cases were reported in 2017 [2]. In 2017, more new cases were detected (214,783 new cases) than those registered for chemotherapy showing that there are some leprosy patients who missed treatment. Unlike registered prevalence, new grade-2 disability cases have only reduced from 14,403 cases in 2007 to 12,819 cases in 2017 [2] which underlines the ineffectiveness of leprosy control strategy. Although the prevalence of leprosy is declining, the stagnant level of new cases detection every year, leprosy induced disabilities, social discrimination and stigma continue to be an important global challenge [3, 4].

Ethiopia is among one of the 122 leprosy endemic countries. The registered prevalence of leprosy in Ethiopia has been reduced from in 5,371 cases in 2007 to 3,692 cases in 2017 which is about 35% reduction in 10 years [2]. On the other hand, new cases detection reduced from 4,187 cases to 3,692 in the same year with a reduction rate of only 6% in ten years. On the contrary the number of patients with G2D has increased from 411 cases in 2007 to 419 cases in 2017 [2]. In the past ten years the proportion of children with grade 2-disabilitties reported in Ethiopia is about 10%. Among the 419 new cases with grade2-disabilities reported in Ethiopia in 2017, 39 of them were children [2]. The high rate of new case detection with high proportion of children with grade 2-disabilities in Ethiopia indicates the presence of active transmission and the late diagnosis of the disease.

In Ethiopia, the integration of leprosy into the general health program has created both challenge and opportunity for effective management of persons affected by leprosy particularly those with disability [5]. It is an opportunity because of the access of health services to those in need which is not possible with the vertical set up. The challenge is that the integration has made that patients are seen by general health workers during outpatient visits that does not have adequate training in leprosy which could lead to misdiagnosis and late treatment [6]. Misdiagnosis and late retreatment are one of the main causes of disability in leprosy patients. Previous study has shown that among 601 general health workers from 300 health centres and 52 general hospitals in Ethiopian only 18% of them correctly diagnose and treat leprosy patients at general public health facilities [5]. Hence, the integration of leprosy into the general health program need building the capacity of health staff for accurate diagnosis and timely treatment of patients.

With the fall in the number of new cases, there is an increasing shift in the focus to prevention of disability [7]. Self-care practice in leprosy is a professionally supported set of practices by leprosy patients to improve and restore health and self-esteem [4, 8]. It is a care that is initiated and maintained by the leprosy patient which needs active engagement of leprosy patient. Hence, Self-care is a process that permeates life and is therefore self-motivated and essentially depends on the commitment of the individual [7]. Proper self-care practice help to reduce leprosy induced disability and stigma. However, the knowledge and perception of leprosy patient could affect the outcome self-care practice [9, 10]. Therefore, understanding the knowledge and perception of leprosy patients towards leprosy self-care practice is critical to develop a meaningful leprosy control and rehabilitation program in the country [11]. Thus, this study was initiated to investigate the knowledge and perception of leprosy patients towards self-care practice at ALERT Hospital, Ethiopia.

## Materials and Methods

### Study design

A cross sectional study was conducted using quantitative methods. The study was done within the framework of leprosy self-care and rehabilitation program at ALERT Hospital.

#### Source of samples and sampling method

Study subjects were obtained from leprosy outpatients attending physiotherapy, red medical clinic, orthopaedic workshop and the ulcer clinic. Patients registered as leprosy case before 2 months to the hospital were considered as a sampling frame. Leprosy patients below 15 years of age and those who did not give their consent were excluded from the study. A systematic random sampling method was used to enrol study subjects to the study. A total of 424 leprosy patients were enrolled to the study.

#### Data collection

Data was collected on structured questionnaire through face to face interview. Core points such as demographic data, early signs and symptoms of leprosy, cause and treatment of leprosy, infectivity, curability and deformity of leprosy. Self-care particle questions such as wound soaking, paraffin lubricant application to the wound, trimming of the wound edge, wound dressing and regular wound cleaning was included to the questionnaires. Perception to the importance of self-care practices such as the importance of soaking, wound trimming, wound cleaning and confidence to the treatment have been included to the questionnaire. The questionnaire was first prepared in English and then translated in to Amharic language. The questioner was administered by eight trained data collectors. Data collectors and 2 supervisors were trained for one day for to create common understanding on each questionnaire. Following the training a pilot test was conducted to validate the data collection tool.

### Operational definitions

We used Bloom’s cut off point to measure knowledge and self-care practice of the respondents [12]. Percent of correct response to a set of 15 knowledge questions and 15 self-care practice questions were used for grading knowledge and self-care practices as follow as:

**Poor knowledge:** Study participants who answered “yes” to 59 % or below (≤8/15) of knowledge and perception questions were considered as having as poor knowledge of leprosy
**Good Knowledge**: Study participants who answered “yes” to above 59 % (>8/15) of knowledge and perception questions were considered as having as Good knowledge of leprosy
**Poor self-care practice**: Study participants who answered “yes” to 59 % or below (≤8/15) of self-care practice questions were considered as having as poor self-care practice
**Good self-care practice**: Study participants who answered “yes” to above 59 % (>8/15) of self-practice questions were considered as having as Good self-practice.

### Statistical analysis

The anonymous demographic and clinical data were entered into an Excel database and analysed using SPSS 23 version-1 statistical software. Chi-square test was used test the level of significance of each variable at α = 5% with 95% confidence interval. Parametric method was used to analyse numerical variables and mean is used for reporting for these variables whenever required.

### Ethical considerations

The protocol was approved by the AHRI/ALERT Ethical Review Committee with approval number of P019/16. The information sheet was read to the participants. Support letter was obtained from the ALERT Hospital. A verbal consent was obtained from each of the participants as recommended by the ethical review committee. Each participant was asked his/her willingness to take part in the survey by the data collector and both the data collector and participant signed on the logbook specifically designed for the survey to keep the list of participants and the data collector name. An assent was obtained for study participants of 15-17 years. Children under 15 years were excluded from the study. All personal information was kept confidential and reporting was made anonymous.

## Results

### Socio-demographic characteristic of study subjects

A total of 424 leprosy patients were requited to the study. The median age of the respondents were 48 years (range: 15-85 years). Among the respondents 236 (55.7%) of them were males. About half of the participants, 226 (53.3%) were married. Considering their employment status 387 (91.3%) of them were employed although majority of them (89.8%) earn less than 1000 ETB/month which is about $1/day (table 1).

**Table 1.** Socio demographic characteristics of the respondents at enrolment.

| Characteristics (Variables) | Number(N) | Percent (%) |
| --- | --- | --- |
| **Age (years)** |  |  |
| 14-23 | 40 | 9.4 |
| 24-33 | 49 | 11.6 |
| 34-43 | 67 | 15.8 |
| 44-53 | 98 | 23.1 |
| > 53 | 170 | 40.1 |
| **Gender** |  |  |
| Male | 236 | 55.7 |
| Female | 188 | 44.3 |
| **Marital status** |  |  |
| Single | 99 | 23.3 |
| Married | 226 | 53.3 |
| Divorced | 52 | 12.3 |
| Separated | 9 | 2.1 |
| Widowed | 38 | 9.0 |
| **Occupational status** |  |  |
| Employed | 387 | 91.3 |
| Unemployed | 37 | 8.7 |
| **Educational status** |  |  |
| Illiterate | 178 | 42 |
| Read and write | 58 | 13.7 |
| Primary | 94 | 22.2 |
| Secondary and above | 94 | 22.2 |
| **Income (ETB)\*** |  |  |
| 0-150 | 116 | 27.4 |
| 151-450 | 109 | 25.7 |
| 451-600 | 73 | 17.2 |
| 601-1000 | 80 | 18.9 |
| > 1000 | 46 | 10.8 |

\* ETB = Ethiopian Birr; Total sample size=424 respondents

### Self-care practice of leprosy patients, ALERT Hospital

Among the 424 respondents only 43 (10.1%) of them use paraffin for lubricating the wound as prescribed by the leprosy care professionals. About one in four patients practice trimming of the wound and only one in ten patients clean the wound. Among the respondents, more than 80% of them do not believe in the importance of wound soaking. On the other hand, more than half of the patients undertake physiotherapy. While nearly half of the patients usually use antiseptic solutions, a third of them practice wood dressing (Table 2).

**Table 2.** Self-care practice of leprosy patients, ALERT Hospital.

| Characteristics | Number | Percent (%) |
| --- | --- | --- |
| Use paraffin lubricant as prescribed |  |  |
| No | 381 | 89.9 |
| Yes | 43 | 10.1 |
| Exercise Trimming of the wound edge |  |  |
| No | 325 | 76.7 |
| yes | 96 | 22.6 |
| Occasionally | 3 | 0.7 |
| Believe Soaking is important |  |  |
| No | 345 | 81.4 |
| Yes | 79 | 18.6 |
| Perform Physiotherapy |  |  |
| No | 198 | 46.7 |
| Yes | 226 | 53.3 |
| Make wound dressing |  |  |
| No | 279 | 65.8 |
| Yes | 145 | 34.2 |
| Clean the wound |  |  |
| No | 395 | 93.2 |
| Yes | 29 | 6.8 |
| Use antiseptic solution usually |  |  |
| No | 223 | 52.6 |
| Yes | 197 | 46.5 |
| Occasionally | 4 | 0.9 |

Total sample size=424 respondents

### Knowledge of the respondents

The knowledge score of the respondent was poor for 276 (65.1%) and good for 148 (34.9%). The level of knowledge was found significantly varied within the age groups (P= 000). As the age of the respondents increase, their knowledge score decrease. Younger age groups (≤ 33 years) had significantly higher knowledge score while older age groups (> 53 years) had the lowest knowledge score. Knowledge score were also significantly different between male and female. While 43.2% male had good knowledge about leprosy only 24.5% of females had good knowledge (P=0.000). Regarding the marital status, single respondents had better leprosy knowledge than the other groups (P-0.003). Only 4.5% respondents who do not write and read had good knowledge score. On the other hand, 63.8% the respondents who are educated to secondary school or above had good knowledge of leprosy compared to the other groups (P=0.000). Unmarried people had good knowledge than people with other marital status (P=0.003). Poor knowledge is a characteristic of peoples who earn less than 150 ETB/month (~$5/month). Access to media and having awareness to leprosy was associated with having good knowledge. On the other hand, there is no statistically significant difference between those who received health education and those who did not receive health education (P>0.05). Similarly, religion and ethnicity had no impact on the level of the knowledge of the respondents (Table 3).

**Table 3.** Knowledge level of leprosy patients towards the disease by socio-demographic characteristics and income status at ALERT Hospital.

| Variable | Poor Knowledge | Good knowledge | Total (N) | Significance test* |
| --- | --- | --- | --- | --- |
| Age (years) | N(%) | N(%) |  |  |
| 14-23 | 15(37.5) | 25(62.5) | 40 | X<sup>2</sup>=22.37<br>p value =0.000 |
| 24-33 | 26(53.1) | 23(46.9) | 49 |  |
| 34-43 | 42(62.7) | 25(37.3) | 67 |  |
| 44-53 | 70(71.4) | 28(28.6) | 98 |  |
| >53 | 123(72.4) | 47(27.6) | 170 |  |
| **Total** | **276 (65.1)** | **148 (34.9)** | **424** |  |
| Sex |  |  |  | X<sup>2</sup>=16.195<br>p value =0.000 |
| Male | 134(56.8) | 102(43.2) | 236 |  |
| Female | 142(75.5) | 47(24.5) | 188 |  |
| **Total** | **276 (65.1)** | **148 (34.9)** | **424** |  |
| Marital status |  |  |  | X<sup>2</sup> =15.932<br>p value = 0.003 |
| Single | 51(51.5) | 48(48.5) | 99 |  |
| Married | 160(70.8) | 66(29.9) | 226 |  |
| Divorced | 28(53.8) | 24(46.2) | 52 |  |
| Separated | 6 (66.6) | 3(33.4) | 9 |  |
| Widowed | 31(81.6) | 7(18.4) | 38 |  |
| **Total** | **276 (65.1)** | **148 (34.9)** | **424** |  |
| Educational status |  |  |  | X<sup>2</sup>=79.185<br>p value =0.000 |
| Illiterate | 170 (95.5) | 08(4.5) | 178 |  |
| Read and write | 28(48.3) | 30(51.7) | 58 |  |
| Primary | 44(46.8) | 50(53.2) | 94 |  |
| Secondary & above | 34(36.2) | 60(63.80) | 94 |  |
| **Total** | **276 (65.1)** | **148 (34.9)** |  |  |
| Income (ETB) |  |  |  | X<sup>2</sup>=15.05<br>p value= 0.005 |
| 0-150 | 95(81.9) | 21(18.1) | 116 |  |
| 151-450 | 84(71.1) | 25(22.9) | 109 |  |
| 451-600 | 49(67.1) | 24(32.9) | 73 |  |
| 601-1000 | 38(47.5) | 42(52.5) | 80 |  |
| > 3000 | 10(21.7) | 36(78.3) | 46 |  |
| **Total** | **276 (65.1)** | **148 (34.9)** | **424** |  |
| Religion |  |  |  |  |
| Orthodox | 227(81.5) | 113(18.5) | 340 | X<sup>2</sup>=3.148<br>p value =0.533 |
| Muslim | 29(60.4) | 19(39.6) | 48 |  |
| Protestant | 17(58.6) | 12(41.4) | 29 |  |
| Catholic | 2(50.0) | 2(50.0) | 4 |  |
| Others | 1(33.3) | 2(66.7) | 3 |  |
| **Total** | **276 (65.1)** | **148 (34.9)** | **424** |  |
| Ethnicity |  |  |  |  |
| Amhara | 186(67.1) | 91(32.9) | 277 | X<sup>2</sup>=8.596<br>p value= 0.072 |
| Oromo | 51(65.4) | 27(34.6) | 78 |  |
| Tigræ | 7(43.8) | 9(56.2) | 16 |  |
| Guragie | 24(70.6) | 10(29.4) | 34 |  |
| Somale | 8(42.1) | 11(57.9) | 19 |  |
| **Total** | **276 (65.1)** | **148 (34.9)** | **148** |  |
| Access of media |  |  |  |  |
| yes | 171(60.4) | 112 (39.6) | 283 | X<sup>2</sup>=5.908<br>p value= 0.015 |
| no | 105(74.5) | 36 (25.5) | 141 |  |
| **Total** | **276 (65.1)** | **148 (34.9)** | **424** |  |
| Awareness of leprosy |  |  |  |  |
| yes | 178(62.9) | 105(37.1) | 283 | X<sup>2</sup>=1.808<br>p value =0.179 |
| no | 98(69.5) | 43(30.1) | 141 |  |
| **Total** | **276 (65.1)** | **148 (34.9)** | **424** |  |
| Confident that they are properly treated |  |  |  |  |
| yes | 215(62.2) | 125(36.8) | 340 | X<sup>2</sup>=4.727 p values =0.030 |
| no | 61(72.6) | 23(527.4) | 84 |  |
| **Total** | **276 (65.1)** | **148 (34.9)** | **424** |  |
| Received health Education |  |  |  |  |
| yes | 228(64.4) | 126(35.6) | 354 | X<sup>2</sup>= 0.446 p value= 0.504 |
| no | 48(68.6) | 22(31.4) | 70 |  |
| **Total** | **276 (65.1)** | **148 (34.9)** | **424** |  |

\* P-value statistically significant at $\alpha=0.05$ ; X<sup>2</sup> = Chi-square; ETB= Ethiopian Birr

### Self care practice

The level of self-care practice was assessed among 424 leprosy patients. Respondents in the age range of 24-33 years had the highest level of self-care practice (44.9%) followed by the age groups 14-23 years (P=0.002). The assessment of self-care practice has shown that as age increases, the level of self-care practice decreases showing that age is an important factor in leprosy self care-practice. Patients who had previous disability due to leprosy had the highest level of self-care practice (62.5%) than those who had no previous disability due to leprosy. Similarly, patients those who had good knowledge (62.8%) about leprosy had good self care practice than those who has poor knowledge (P= 0.038). With regard to the income status of the respondents, as income increases, the level of self-care practice increases (table 4).

Although sex is found to be associated with knowledge that males had more leprosy knowledge compared to females, the level of self-care practice to leprosy is not significantly different between males and females. Similarly, educational and marital status of the respondents did not associated with leprosy self-care practice (P>0.05). Interestingly, leprosy self-care practice was not significantly different between those who received health education and those who did not receive health education (table 4).

**Table 4.** Self care practice of leprosy patients at ALERT Hospital.

| Variable | Self care practice |  | Total Number (N) | significance level * |
| --- | --- | --- | --- | --- |
|  | Poor | Good |  |  |
| **Age (years)** | N(%) | N(%) | | $X^2= 17.001$<br>p value= 0.002 |
| 14-23 | 30(75.0.) | 10(25.0) | 40 |  |
| 24-33 | 27(55.1) | 22(44.9) | 49 |  |
| 34-43 | 52(77.6) | 15(22.4) | 67 |  |
| 44-53 | 79(80.6) | 19(19.4) | 98 |  |
| >53 | 140(82.4) | 30(17.6) | 170 |  |
| Total | 328 (77.4) | 96(22.6) | 424 |  |
| **Sex** | | | | $X^2= 0.643$<br>p value= 0.422 |
| Male | 50(21.2) | 186(78.8) | 236 |  |
| Female | 46(25.5) | 142(74.5) | 188 |  |
| Total | 96(22.6) | 328 (77.4) | 424 |  |
| **Previous disability due to leprosy** |  |  |  |  |
| Yes | 123(37.5) | 205(62.5) | 328 | $X^2 =49.241$<br>p value =0.000 |
| No | 75(78.1) | 21(21.9) | 96 |  |
| Total | 198 (46.7) | 226 (53.3) | 424 |  |
| **Knowledge of leprosy** | | | | $X^2 =4.291$<br>p value =0.038 |
| Poor knowledge | 204(73.9) | 72(26.1) | 276 |  |
| Good knowledge | 55(37.2) | 93(62.8) | 148 |  |
| Total | 259 (61.1) | 165 (38.9) | 424 |  |
| **Educational status** |  |  |  |  |
| Illiterate | 140(78.7) | 38(21.3) | 178 | $X^2=5.069$<br>p value =0.167 |
| Read and write | 48(82.8) | 10(17.2) | 58 |  |
| Primary | 75(79.8) | 19(20.2) | 94 |  |
| Secondary and above | 65(69.1) | 29(30.9) | 94 |  |
| Total | 328(77.4) | 96 (22.6) | 424 |  |
| **Income (ETB)** |  |  |  |  |
| 0-150 | 97(83.6) | 19(16.4) | 116 | X<sup>2</sup>=10.853<br>p value= 0.028 |
| 151-450 | 90(82.6) | 19(17.4) | 109 |  |
| 451-600 | 59(80.8) | 14(19.2) | 73 |  |
| 601-1000 | 55(68.8) | 25(31.3) | 80 |  |
| >1000 | 27(58.7) | 19(41.3) | 46 |  |
| Total | 328(77.4) | 96 (22.6) | 424 |  |
| **Marital status** |  |  |  |  |
| Single | 72(72.7) | 27(27.3) | 99 | X<sup>2</sup>= 4.537<br>p value = 0.338 |
| Married | 173(76.5) | 53(23.5) | 226 |  |
| Divorced | 45(86.5) | 7(13.5) | 52 |  |
| Widowed/widower | 8(88.9) | 1(11.1) | 9 |  |
| Separated | 30(78.9) | 8(21.1) | 38 |  |
| Total | 328(77.4) | 96 (22.6) | 424 |  |
| **Received Health Education** |  |  |  |  |
| Yes | 277(78.2) | 77(21.6) | 354 | X<sup>2</sup>=0.970<br>p value =0.325 |
| No | 51(72.9) | 19(27.1) | 70 |  |
| Total | 328(77.4) | 96 (22.6) | 424 |  |

\* P-value statistically significant at $\alpha=0.05$ ; X<sup>2</sup> = Chi-square; ETB= Ethiopian Birr

## Discussion

As the study reveals nine in 10 peoples earn less than $1/day which indicates the possibility of the association of leprosy and poverty in Ethiopia. Although it is very difficult to demonstrate the link between poverty and leprosy at community or individual level, several epidemiological studies have consistently reported the strong association between leprosy and poverty at community and household level in Indonesia [13] in Bangladesh [14], in Malawi [15] and in India [16].

Generally it was found that leprosy patients had poor knowledge about the disease and poor self-care practice. Over the past two decades leprosy program has initiated the integration of leprosy control services into the general health care services in Ethiopia [5]. However, it has been reported that the integration has made that patients are seen by general health workers during outpatient visits rather than by leprosy specialized personnel in leprosy dedicated clinics which could lead to misdiagnosis and treatment [5]. It has been a concern that the integration has given inadequate attention towards activities of prevention of disabilities and rehabilitation in the integration process [17].

In leprosy, self-care is the most important type of self-management where leprosy patients need to change their behavior to adapt to the irreversible impairment due to the disease[3]. Although, limited information is available on the effectiveness of self care management, it has been reported that self-care practice particularly participation of patients in self-care group improves their understanding about self-care. Self-care practice helps patients to reduce the number of ulcers, improve their physical condition and increase self-confidence [3]. However, the effectiveness of self-care practice can be compromised by poor knowledge and perception about the disease. In our study we found that poor self care-practice was associated with age, sex, marital status, educational status, income and access to media.

Age of the patients was one of the variable which significantly associated to leprosy knowledge and self-care practice (table 2 and 3). Patients in the age range of 14-23 years had significantly higher knowledge and good self-care practice compared to the other age groups. This could be due to the fact that people in this age group are more likely to take care of themselves as they are in the peer-forming e age group. Several studies have reported that leprosy influences marriage and sexual relationship in several countries such as Nepal, India, Bangladesh and Brazil [18, 19]. In our study, we found that females had poor leprosy knowledge and poor self –care practice to leprosy compared to males. Like in many developing countries, in Ethiopia women play a subordinate, submissive and more conservative gender role in seeking health care particularly in poverty stricken rural and remote areas of the country [20]. Societal stigma, women’s dependence and low status, self-stigmatizing attitudes, and the gender insensitivity of leprosy services are reported as barriers faced by women [21]. Hence, more work is yet being to be done by health care professionals and the society to tackle these barriers in Ethiopia where leprosy is endemic.

Being single is associated with relatively improved self-care and having good comparative knowledge of leprosy compared to the other marital status. This could be due to the fact that singles are more concerned about their future relationship and they want to recover from leprosy as quickly as possible. Patients with no educational history and those who have no access to any form of media had less knowledge of leprosy and had poor self-care practice. Several studies have reported that educated leprosy patients have good leprosy knowledge and leprosy self-care practice compared to uneducated patients [22–24]. Hence, leprosy education and rehabilitation program may need adult literacy program combined with more innovative focused approaches to suit various target audiences that can impact knowledge, attitudes and self-care practice better.

In conclusion, this study has identified that majority of leprosy patients (65%) have poor knowledge and poor leprosy self-care practices in spite of the fact that leprosy treatment, disability prevention and rehabilitation program run in the country under general public health care service for decades. The leprosy service program should re-visit its strategy and mode of delivery to improve the leprosy knowledge and leprosy self-care practices of patients.

## Supporting information legend

S1 Checklist STROBE Checklist

